# Aging is associated with increased brain iron through brain-derived hepcidin expression

**DOI:** 10.1101/2021.09.06.459092

**Authors:** Tatsuya Sato, Jason Shapiro, Hsiang-Chun Chang, Richard A. Miller, Hossein Ardehali

## Abstract

Iron is an essential molecule for biological processes, but its accumulation can lead to oxidative stress and cellular death. Due to its oxidative effects, iron accumulation is implicated in the process of aging and neurodegenerative diseases. However, the mechanism for this increase in iron with aging, and whether this increase is localized to specific cellular compartment(s), are not known. Here, we measured the levels of iron in different tissues of aged mice, and demonstrate that while cytosolic non-heme iron is increased in the liver and muscle tissue, only the aged brain exhibits an increase in both the cytosolic and mitochondrial non-heme iron. This increase in brain iron is associated with elevated levels of local hepcidin mRNA and protein in the brain. We also demonstrate that the increase in hepcidin is associated with increased ubiquitination and reduced levels of the only iron exporter, feroportin-1 (FPN1). Overall, our studies provide a potential mechanism for iron accumulation in the brain through increased local expression of hepcidin, and subsequent iron accumulation due to decreased iron export. Additionally, our data support that aging is associated with mitochondrial and cytosolic iron accumulation only in the brain and not in other tissues.

## Introduction

Iron is an essential molecule for almost every organism on earth. It is generally found naturally in two oxidation states, either as the oxidized form of ferric iron (Fe^3+^), or as the reduced form of ferrous iron (Fe^2+^). Ferrous iron is the major form of iron used in mammalian cells, and due to the scarcity of the reduced form of iron on earth, mammalian cells have developed mechanisms for efficient uptake of oxidized iron through its solubilization by acidification, followed by reduction and its cellular transportation. Iron can donate and accept electrons from various substrates due to its unique oxidation-reduction properties, making it an important cofactor in mammalian cells. Iron is also essential for heme and Fe-S clusters and exerts other biological effects through its role in processes such as demethylation, dehydrogenation and reduction of sulfur (Koleini et al., 2021).

Due to its oxidative effect, iron accumulation is hypothesized to lead to oxidative stress and cellular damage that accelerates the process of aging (Xu et al., 2012). Aging has been shown to be associated with iron accumulation in various organs and tissues both in humans and in animal models of aging (Xu et al., 2012). However, whether iron accumulation occurs in all organs or is specific to one organ is not known. In the brain, iron accumulation has been demonstrated both in animal models of aging (Benkovic and Connor, 1993; Hahn et al., 2009; Roskams and Connor, 1994) and in humans (Bartzokis et al., 2010; Bartzokis et al., 1994a; Bartzokis et al., 1994b; Zecca et al., 2001). Studies using MRI of the brain have also demonstrated increased brain iron, and a meta-analysis of these studies supported an association with aging and increased iron content in the substantia nigra and stratum (Daugherty and Raz, 2013). It has also been reported that brain iron accumulation is associated with neurodegenerative diseases like Alzheimer’s and Parkinson’s disease (Berg et al., 2001; Smith et al., 1997). Additionally, it is now believed that iron accumulation may be the cause of neuronal cell death in some of neurodegenerative diseases (Lei et al., 2012). Since the risk of Alzheimer’s and Parkinson’s disease are both increased with aging, it is possible that iron accumulation due to advanced age may play a major role in the development of age-associated neurodegenerative diseases.

Despite the importance of iron in brain disorders, the mechanism by which iron accumulates in the brain with aging is not known. Previous studies suggest that iron homeostasis in the brain is dependent on normal expression of genes involved in iron uptake into the cells and its cellular storage and regulation (Qian and Wang, 1998). Another study demonstrated that only one protein involved in the iron import pathway, divalent metal transporter (DMT1) was altered in aged brain, while the levels of key proteins involved in iron import and export, transferrin receptor-1 (TfR1) and feroportin-1 (FPN1) were unchanged (Lu et al., 2017). However, the mechanism for the age-related increase in iron accumulation in the brain remains unclear.

In this paper, we assessed the levels of iron in different tissues and cellular compartments with aging in mice. We demonstrate that the cytosolic non-heme iron is increased in the muscle and liver tissue, while the brain has increases non-heme iron both in the mitochondria and in the cytosol. We also demonstrate that hepcidin levels are upregulated in the brain of aged mice, which is associated with ubiquitination of FPN1 and a reduction in its protein levels. Thus, our studies suggest that local expression of hepcidin in the brain may be a driver in increasing iron levels due to a reduction in iron export by FPN1 degradation.

## Results

### Iron is increased in the brain with aging

To determine whether iron accumulates in tissues with aging, we measured heme and non-heme iron levels in the mitochondria and cytoplasm of liver, skeletal muscle and brain tissues. These organs were selected since liver is the main iron storage organ, while skeletal muscle is a major iron consuming organ. Additionally, brain iron accumulation is reported with neurodegenerative diseases. We chose two time points for our studies: 4 month for young mice and 22 month for old mice. The almost 2 year age of the old mice is considered sufficient to result in senescence of some organs (https://www.jax.org/news-and-insights/jax-blog/2017/november/when-are-mice-considered-old#). Our data demonstrated that cytosolic non-heme iron in the liver was increased with age, whereas liver iron contents in the mitochondria, where iron accumulation is associated with cell dysfunction and cell death, were unchanged with age for both heme and non-heme forms (**Figure 1A and B**). Similarly, we observed an increase in the cytosolic, but not mitochondrial, non-heme iron in the gastrocnemius muscle. We also did not see a change in the cytosolic heme iron, however, we were not able to detect heme iron in the mitochondria of skeletal muscle tissue, possibly due to technical limitations associated with the isolation of intact mitochondria from skeletal muscle tissue (**Figure 1C and D**).

**Figure 1.**
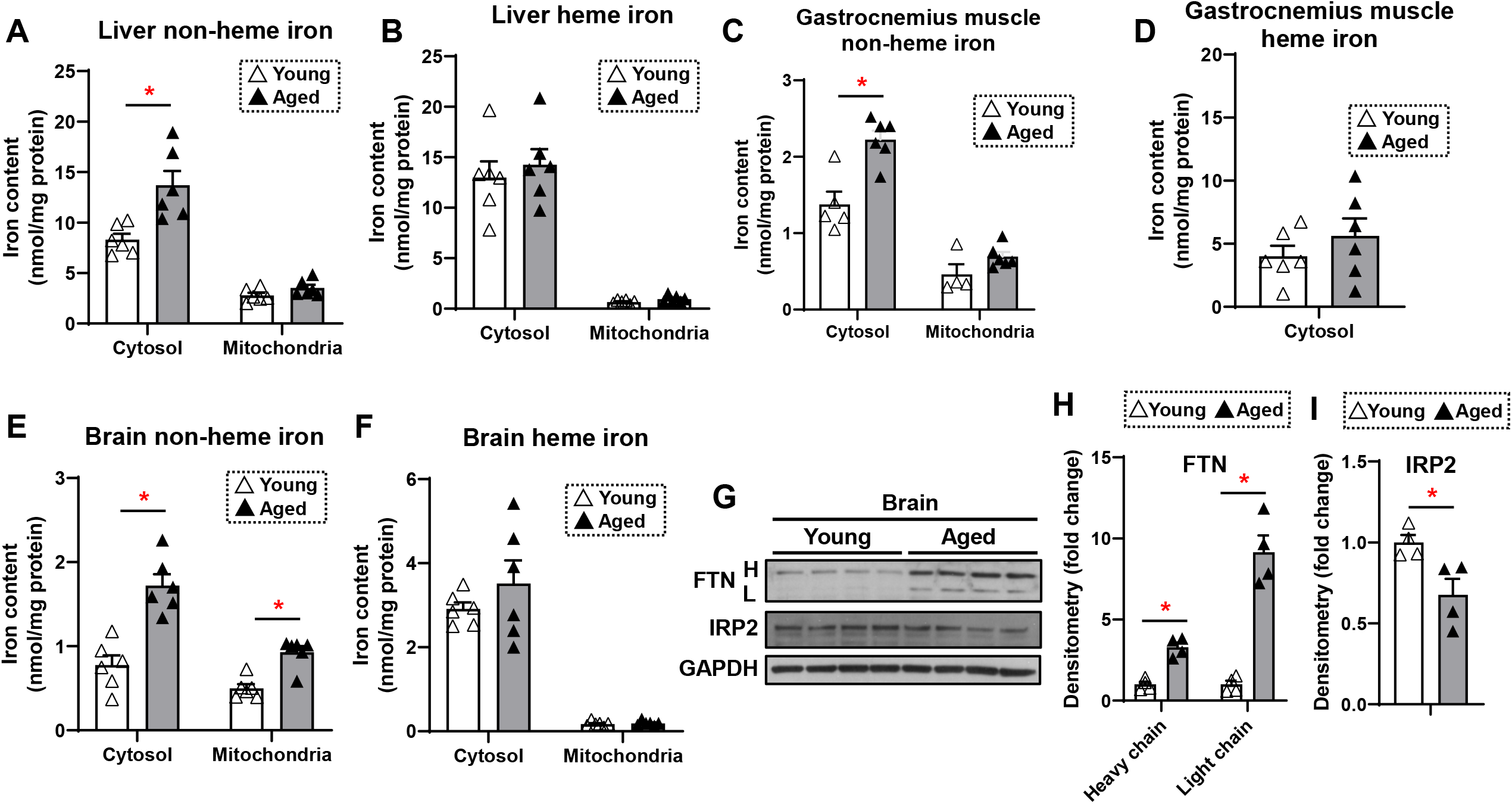
Non-heme iron is increased in the cytosol and mitochondria of the brain cortex with aging. Liver non-heme (**A**) and heme (**B**) iron in young (4 month old) and old (22 month old) mice from the cytosol and mitochondria (n=6). Gastrocnemius non-heme (**C**) and heme (**D**) iron in young and old mice from the cytosol and mitochondria (n=6). Outliers in non-heme iron in the cytosol (n=1) and in the mitochondria (n=2) were excluded. Mitochondrial heme iron in gastrocnemius muscle was not detectable in our studies. Brain non-heme (**E**) and heme (**F**) iron in young and old mice from the cytosol and mitochondria (n=6). (**G**) Immunoblots of FTN and IRP2 in the brain of young and aged mice (n=4). FTN = ferritin, IRP2 = iron regulatory protein 2, H = heavy chain, L = light chain. Densitometric quantification of light and heavy chain FTN and IRP2 in the brain of young and aged mice. ^*^ P< 0.05.

We then studied iron levels in the brain mitochondrial and cytosolic fractions, and noted a significant increase in non-heme iron in both the mitochondria and cytoplasm of brain tissue in older mice (**Figure 1E**). However, there was no increase in heme iron levels in the either compartment in the brain (**Figure 1F**). These results suggest that the more oxidative form of iron (i.e., non-heme iron) is increased in the brain cytosol and mitochondria with aging. Consistent with increased non-heme iron levels, we also observed increased protein levels of ferritin light and heavy chain (**Figure 1G and H**). This increase may be a cellular protective mechanism because free or loosely-bound iron in the form of non-heme iron is detrimental to the cells, and iron stored in ferritin is considered redox-inert. Additionally, an elevated level of cellular iron leads to degradation of IRP2 protein (Wang et al., 2004). We therefore measured IRP2 protein levels in the brain and observed a decrease in IRP2 protein levels in the brain of aged mice (**Figure 1G and H**), consistent with an increase in iron levels with aging.

### Hepcidin levels are increased in the brain tissue with aging

Since non-heme iron is increased in both the cytosolic and mitochondrial compartments of the brain and previous reports suggesting a potential role for iron in neurodegenerative diseases associated with aging (Berg et al., 2001; Smith et al., 1997), we then focused our studies on the mechanism for the increase in iron in the brain tissue. We measured the mRNA levels of all proteins involved in cellular iron homeostasis. We conducted these studies both in the liver and brain tissue. In the liver, we noted a significant increase in the mRNA levels of *Fpn1*, heme oxygenase 1 (*Hox1*), ALA dehydrogenase (*Alad*), and bone morphogenetic protein 6 (*Bmp6)*, and a significant decrease in the mRNA level of *Steap3* (**Figure 2A**). Since none of these changes lead to increased cellular iron levels, we concluded that the alterations of the mRNA levels are not responsible for changes in cellular iron levels. However, we noted a significant increase in the mRNA levels of *TfR2, Mfrn2, Ttp, Hox1, Alad* and hepcidin (*Hamp1)* in the brain tissue of aged mice (**Figure 2B**). TFR2 does not play a meaningful role in iron uptake compared to transferrin-receptor protein (TFRC), since it has a much lower affinity for transferrin (Kawabata et al., 1999). Instead, it acts as a signaling molecule to regulate HAMP1 transcription through SMAD signaling (Silvestri et al., 2014). Since both TFR2 and HAMP1 are increased in the brain with aging, we then focused our studies on the cellular hepcidin pathway. To confirm that the increase in *Hamp1* mRNA with aging is associated with an increase in its protein, we measured HAMP1 protein in the brains of young and aged mice and noted a significant increase in its protein levels (**Figure 2C and D**). Finally, we performed immunohistochemistry on the brain of aged mice and confirmed that HAMP1 protein levels are significantly increased with aging (**Figure 2E**).

**Figure 2.**
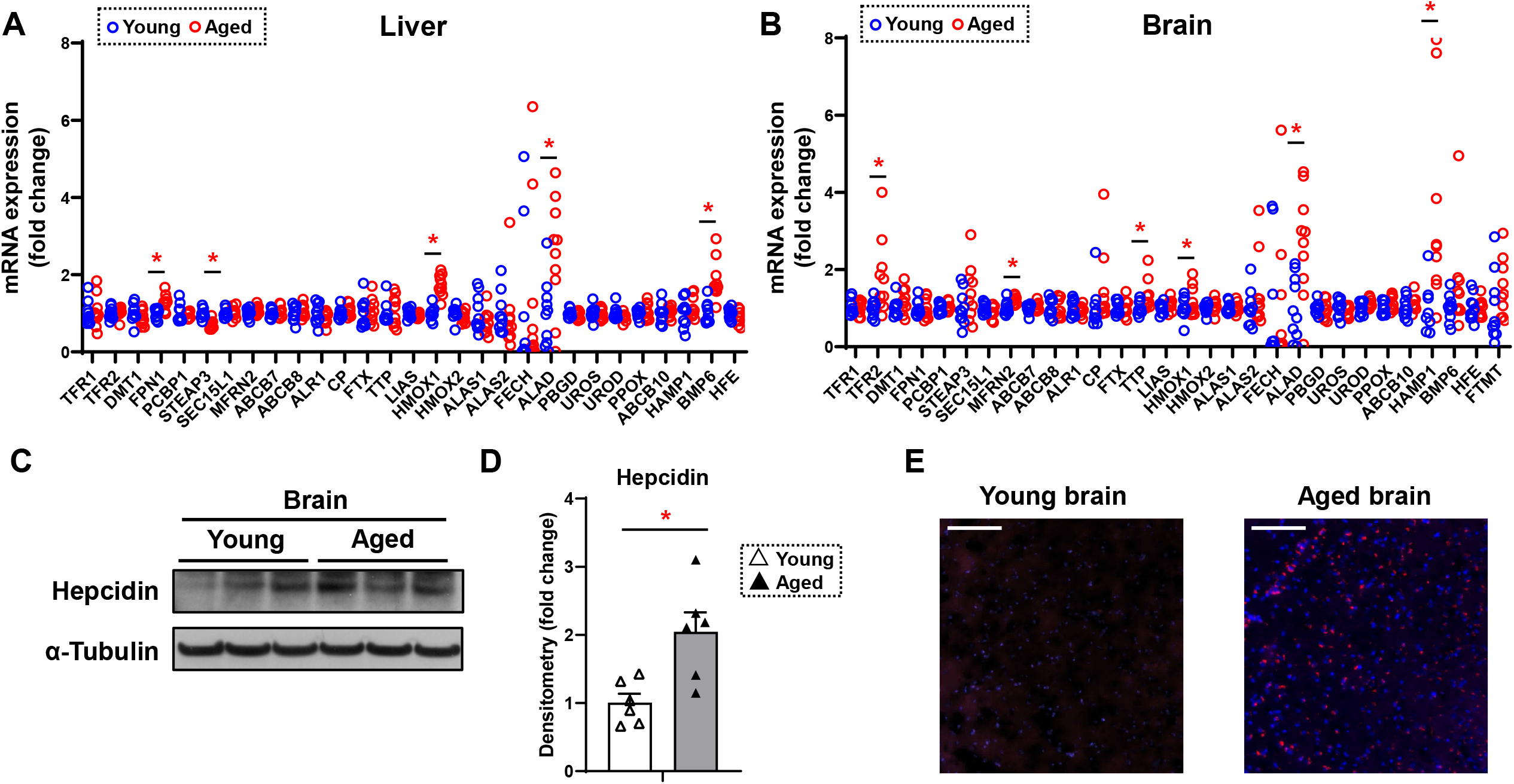
Hepcidin protein expression is significantly increased in the aged brain. mRNA levels of proteins involved in iron regulation in the liver (**A**) and brain (**B**) in young (4 month old) and old (22 month old) mice (n = 10/group). An undetected measurement (n=1) in the brain FECH and undetected measurements (n=2) and an outlier (n=1) in the brain HAMP1 were excluded. TFR1 = transferrin receptor 1, TFR2 = transferrin receptor 2, DMT1 = divalent metal transporter 1, FPN1 = ferroportin1, PCBP1 = Poly(RC) Binding Protein 1, STEAP3 = Metalloreductase STEAP3, Sec15l1 = exocyst complex component 6, MFRN2 = mitoferrin 2, ABCB7 = ATP-binding cassette sub-family B member 7, ABCB8 = ATP-binding cassette sub-family B member 8, ALR1 = Mg(2+) transporter ALR1, CP = ceruloplasmin, FTX = frataxin, HO1 = heme oxygenase 1, HO2 = heme oxygenase 2, ALAS1 = 5’-aminolevulinate synthase 1, ALAS2 = 5’-aminolevulinate synthase 2, FECH = ferrochelatase, ALAD = aminolevulinate dehydratase, PBGD = porphobilinogen deaminase, UROS = uroporphyrinogen III synthase, UROD = uroporphyrinogen decarboxylase, PPOX = protoporphyrinogen oxidase, ABCB10 = ATP-binding cassette, sub-family B member 10, HAMP1 = hepcidin1, BMP6 = bone morphogenetic protein 6, HFE = homeostatic iron regulator, FTMT = mitochondrial ferritin. (**C**) Representative immunoblot for hepcidin1 in the brain (n=6). (**D**) Summary of densitometry analysis of Panel C. (**E**) Representative immunohistochemistry of hepcidin1 (Red = anti-hepcidin1, blue = DAPI) in the brain of young and aged mice. Scale bar = 200 µm. ^*^ P< 0.05.

### Aging is associated with decreased FPN1 protein levels through its ubiquitination

The major effect of hepcidin protein on iron regulation is through ubiquitination and subsequent degradation of FPN1, resulting in a decrease in cellular iron export (Qiao et al., 2012). This mechanism of hepcidin effect generally occurs in intestinal cells and macrophages, which lead to decreased iron export from these cells. Since HAMP1 protein levels are increased in the brain tissue of aged mice, we hypothesized that FPN1 levels would be decreased in the brains of these mice. FPN1 protein levels are significantly decreased in the brain tissue of aged mice, while TfR1 levels are unchanged (**Figure 3A and B**). Since mRNA levels of FPN1 are not significantly changed in the brain, we hypothesized that the decrease in FPN1 protein might be due to its degradation by hepcidin. We therefore measured FPN1 protein ubiquitination in the brains of aged mice, and showed a significant increase in its ubiquitination (**Figure 3C and D**). These results indicate that FPN1 protein is ubiquitinated and decreased with aging, perhaps as a consequence of the higher levels of hepcidin.

**Figure 3.**
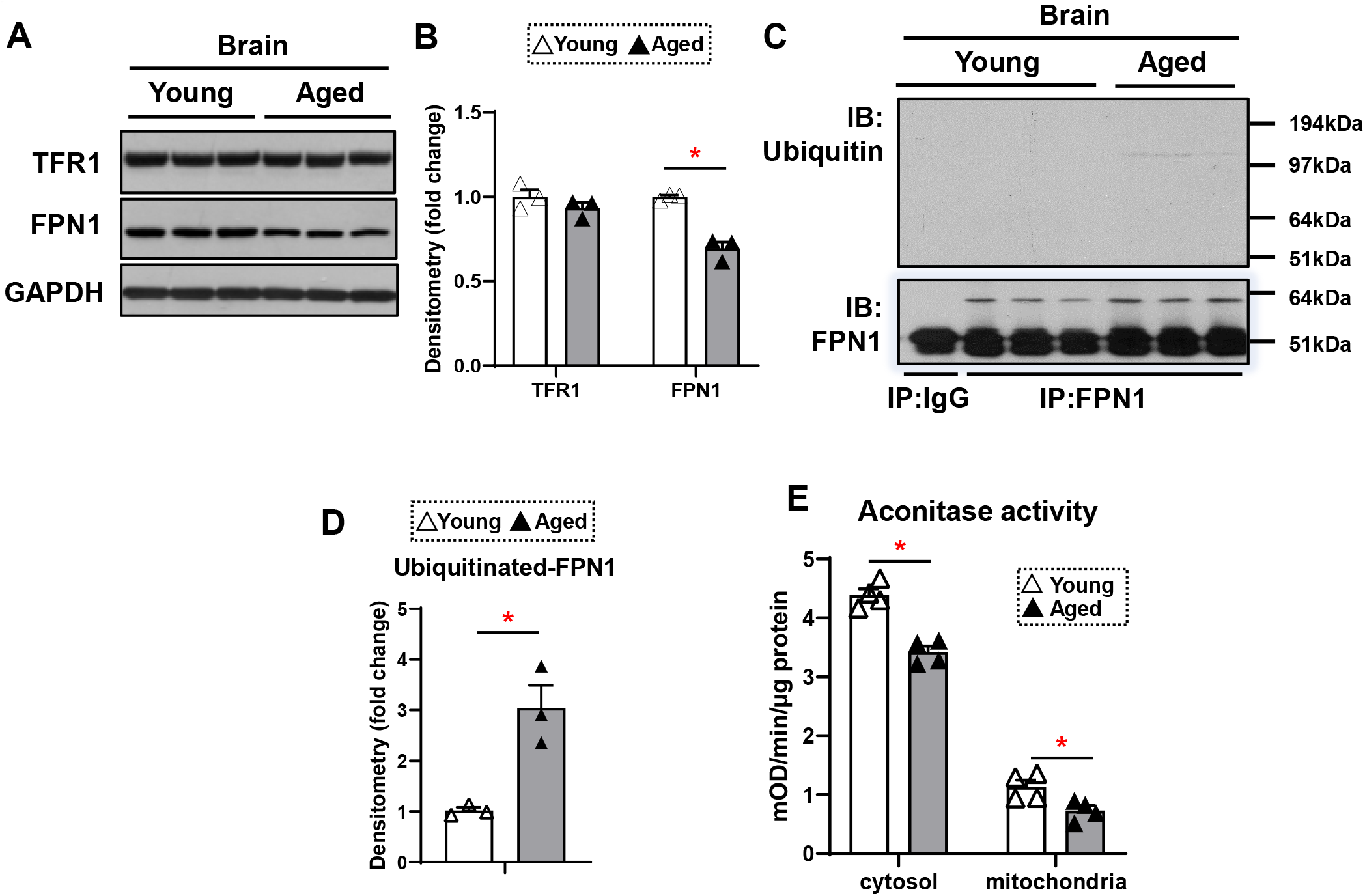
FPN1 protein level is decreased while its poly-ubiquitination is increased in the brain of aged mice. (**A**) Immunoblots of iron transporting proteins TfR1 and FPN1 in the brain of young and aged mice (n=3). (**B**) Summary of densitometric analysis of panel A. (**C**) Poly-ubiquitination levels of FPN1, as assessed by immunoprecipitation, in the brain of young and aged mice (n=3). (**D**) Summary of the densitometric analysis of panel C. (**E**) Fe-S cluster containing aconitase enzyme activity, a marker of cellular oxidative stress, in the brain of young and aged mice (n=4). c-aconitase = cytosolic aconitase (ACO1), m-aconitase = mitochondrial aconitase (ACO2). ^*^ P< 0.05.

We then assessed whether the increase in iron in the brains of aged mice is associated with increased oxidative stress. Aconitase is an Fe/S containing protein that is highly sensitive to oxidative stress (Noster et al., 2019). Our results demonstrated a decrease in both cytosolic and mitochondrial aconitase activity in the brains of aged mice (**Figure 3E**), indicating increased oxidative stress.

## Discussion

Iron is critical for normal cell function, but its excess can lead to cellular damage due to oxidative stress. Thus, the levels of cellular iron, particularly in the mitochondria, are tightly regulated (Koleini et al., 2021). Iron has been implicated in the development of some neurodegenerative diseases that are associated with aging, including Alzheimer’s and Parkinson’s disease (Xu et al., 2012). However, the molecular mechanism for the increase in brain iron in aging is not known. In this paper, we showed that iron accumulates in the mouse brain with aging and that this is associated with a significant increase in hepcidin production in the brain. Since the major function of hepcidin is to bind to FPN1, leading to FPN1 degradation, we assessed levels of FPN1 in brain extracts from young and aged mice and show that while its mRNA is not changed, protein levels of FPN1 are significantly decreased. We also demonstrate that brain FPN1 ubiquitination is increased with aging. Finally, we demonstrate that there is increased oxidative stress in the brains of aged mice, which may well reflect, at least in part, accumulation of iron, perhaps in the context of other age-dependent changes in oxidative damage and defenses against oxidative injury.

Hepcidin antimicrobial peptide (HAMP1) is a small 25–amino acid hormone peptide that is secreted by hepatocytes in conditions of iron sufficiency, and which binds to FPN1 and promotes its internalization and degradation. FPN1 degradation inhibits iron absorption from the digestive tract and the release of iron from macrophages. Although liver is believed to be a major source of systemic hepcidin production, this peptide has recently been shown to be expressed by other tissues. A recent study demonstrated that hepcidin is expressed in the heart and that its cardiac-specific deletion has detrimental effects on cardiac function (Lakhal-Littleton et al., 2016). Additionally, hepcidin has been inferred to play a role in colorectal cancer by sequestering iron to maintain the nucleotide pool and sustain proliferation in colorectal cancer (Schwartz et al., 2021). Recent studies have also demonstrated that hepcidin overexpression in astrocytes protects against amyloid-β induced brain damage in mice and also protects against mouse models of Parkinson’s disease (Liang et al., 2020; Zhang et al., 2020). Our data show that brain hepcidin is increased with aging and that it likely leads to a reduction in FPN-1 levels and iron accumulation. Our studies do not identify the mechanism by which hepcidin is increased with aging, but another study suggested that toll-like receptor 4/myD88-mediated signaling of hepcidin expression cause brain iron accumulation and oxidative injury (Xiong et al., 2016).

The mechanism for iron mediated aging is likely due to the oxidative effects of iron and cellular damage. Mitochondria are the major site of reactive oxygen species production in the cells, thus, the levels of mitochondrial iron are tightly controlled. In our studies, we noted an increase only in the cytosolic non-heme iron in the liver and the skeletal muscle of aged mice, while non-heme iron was increased in both the mitochondrial and cytosolic fraction of the brain tissue. The significance of sole increase in cytosolic iron is not clear, but the increase in mitochondrial iron in the brain is significant since it likely contributes to the oxidative damage that occurs in this tissue with aging. It is also possible that cytosolic iron accumulation in the liver and muscle tissue is due to mild increased systemic iron. Since the brain is not an iron storage organ and iron movement to the brain has to go through the blood-brain barrier (BBB), iron accumulation in the brain with aging is likely independent of systemic iron metabolism.

Our studies have clinical implications for neurodegenerative diseases associated with aging through two possible mechanisms: 1) by targeting brain iron with iron chelators that can cross the blood-brain barrier, and 2) through a reduction in locally produced hepcidin. There is currently interest in taking the first approach and clinical studies support this approach (Sun et al., 2018). However, targeting hepcidin in the brain is technically challenging and may require the design of small molecule hepcidin inhibitors that can also cross the BBB.

## Materials and Methods

### Tissues from Young and Aged Mice

Tissues of liver, gastrocnemius muscle, and brain cortex from young (4 months old) and aged (22 months old) female UM-HET3 mice were used in the present study. For hepcidin immunohistological analysis, young (4 months old) and aged (22 months old) female C57BL/6 mice were used. This study was performed in strict accordance with the recommendations in the Guide for the Care and Use of Laboratory Animals of the National Institutes of Health. All of the animals were handled according to approved institutional animal care and use committee (IACUC) of Northwestern University.

### Cytosolic and Mitochondrial Fractioning

Cytosolic and Mitochondrial fractions from tissues were isolated using Mitochondrial Isolation Kit for Tissues (Pierce) according to manufacturer’s protocol.

### Non-heme and Heme Iron Measurement

Iron contents in the cytosolic and mitochondrial fractions of the tissues were measured as previously described (Bayeva et al., 2012; Sato et al., 2018). Briefly, for the non-heme iron measurements, equal amounts of protein were mixed with protein precipitation solution (1:1 1N HCl and 10% trichloroacetic acid) and heated at 95°C for 1 hour to release iron. The precipitated proteins were removed by centrifugation at 16,000 g for 10 minutes at 4°C, the supernatant was mixed with the equal volume of chromogenic solution (0.5 mM ferrozine, 1.5 M sodium acetate, 0.1% [v/v] thioglycolic acid), and the absorbance was measured at 562 nm. For the heme iron measurements, equal amounts of protein were mixed with 2.0 M oxalic acid and were heated at 95°C for 1 hour to release iron from heme and generate protoporphyrin IX. Samples were then centrifuged at 1,000 g for 10 minutes at 4°C and the fluorescence of the supernatant was assessed at 405 nm/600 nm.

### Reverse Transcription and Quantitative Realtime PCR

RNA was isolated from tissues using RNA-STAT60 (Teltest). Reverse transcribed with qScript Reverse Transcription Kit (Quanta) according to manufacturers’ instruction, and as described previously (Chang et al., 2021). The resulting cDNA was amplified quantitatively using PerfeCTa SYBR Green Mix (Quanta) on a 7500 Fast Real-time PCR System (Applied Biosystems). The relative gene expression was determined using differences in Ct values between gene of interest and house-keeping control genes β-actin and HPRT1.

### Immunoblotting

Tissues were lysed in radioimmunoprecipitation assay (RIPA) buffer supplemented with protease inhibitor (Fisher Scientific). Protein concentration in lysates was determined using BCA Protein Quantification Kit (Pierce) and as described previously (Sawicki et al., 2018). Equal amounts of proteins were resolved on 4-12% Novex Bis-Tris poly-acrylamide gel (Invitrogen) and blotted onto nitrocellulose membrane (Invitrogen). After blocking with tris-buffered saline containing 0.05% Tween 20 (Fisher) and 5% milk, the membrane was incubated overnight at 4°C in primary antibody against FTN (Sigma), IRP2 (Novus Biologicals), TFR1 (Invitrogen), FPN1 (Novus Biologicals), Hepcidin (Abcam), Ubiquitin, GAPDH (Santa Cruz), and α-Tubulin (Abcam).The following day, membranes were incubated with HRP-conjugated anti-mouse or anti-rabbit secondary antibodies (Santa Cruz) for 1 hour and proteins were visualized using Super Surgical Western Pico ECL substrate (Pierce). Quantification of immunoblotting image was done using ImageJ (NIH).

### Immunoprecipitation of FPN1

Tissues were lysed in IP buffer containing 25 mM Tris-HCl (pH 7.5), 150 mM NaCl, 1 mM EDTA, 0.1% NP-40 with protease inhibitor cocktail (Fisher Scientific). 400μg of protein was pre-incubated with 40μl of protein G magnetic beads (Invitrogen) for 1 hour at 4°C. After the beads had been discarded, the supernatant was incubated with FPN antibody (Novus Biologicals) or IgG at a rotator overnight at 4°C.The mixture was then incubated with 40μl of fresh beads for 1 hour at 4°C. A magnetic field was applied to this mixture, and the supernatant was removed. The magnetic beads were washed 3 times with 400μl of IP buffer, re-suspended in 40μl of SDS sample buffer, and incubated for 10 minutes at 70 °C. Finally, 20μl of the supernatant was collected after applying a magnetic field to the mixture and was used for immunoblotting.

### Immunohistochemistry of hepcidin

Mice were anesthetized with an intraperitoneal injection of 250 mg/kg dose of freshly prepared Tribromoethanol (Avertin). They were then transcardially perfused with ice-cold phosphate buffered saline to wash out the blood, and following the discoloration of liver, the buffer was replaced to freshly prepared ice-cold 4% paraformaldehyde (PFA) with the assistance of circulating pump to supply sufficient perfusion pressure. After 20 minutes perfusion, the brain tissue was extracted from the skull and was fixed overnight in 4% PFA.The fixed brain tissue was placed in 30% sucrose for 48 hours at 4°C. Brain tissue was then submerged into OTC compound (Sakura Finetek, Torrance, CA) and frozen in liquid nitrogen. 30µm coronal sections were cut using a cryotome (Leica) and mounted on Fisherbrand™ Superfrost™ Plus Microscope Slides. Sections were permeabilized using 0.25% Triton-x100 in PBS for 30minutes, washed 3x in PBS, and incubated with primary antibody against hepcidin overnight at 4ºC. Sections were then washed 3 times in PBS and incubated with Alexa Fluor®594 Goat Anti-Rabbit IgG at 1:200 (Jackson Immunoresearch) for 2 hours at room temperature. Nuclei were counterstained with DAPI containing ProLong™ Gold Antifade mounting media (Invitrogen) and images were acquired using a Zeiss Axio Observer.Z1 fluorescence microscope.

### Aconitase activity measurements

Aconitase activities in cytosolic and mitochondrial fraction of the brain cortex lysates were measured using Aconitase Enzyme Activity Microplate Assay Kit (Abcam) according to manufacturer’s instructions.

### Statistical Analysis

Data are presented as mean ± SEM. There was no randomization of the samples, but the investigator performing the measurements was blinded to the group assignment. For sample size estimation, our previous study (Chang et al., 2016)) indicated that changes in mitochondrial non-heme iron contents of about 20% can affect iron-dependent cytotoxicity in the mouse heart. Given an estimated standard deviation of iron of 15% in each group, we estimated that 6 samples would be needed to detect a 20% difference in change of iron contents. For other experiments, power analysis was not performed a priori because we could use sufficient numbers of stored tissue samples that had already been cryopreserved and there was no risk of wasting unnecessary animals. We determined n=10 for the RNA levels and n=3∼6 for the immunoblots and enzyme activities for our analysis. This was based on our prior experience performing similar experiments. Unpaired two-tailed Student’s t tests were used to determine statistical significance. P< 0.05 was considered to be statistically significant, as indicated by an asterisk. Analysis was performed using Graphpad Prism 9.

## Acknowledgements

We would like to thank technical help from Chunlei Chen. T.S. was supported by American Heart Association. H.A. is supported by NIH R01 HL127646, R01 HL140973, and R01 HL138982. R.A.M. was supported by NIH grants U01-AG022303-17 and P30-AG024824.

## Author Contributions

T.S. and H.A. designed and performed the research and wrote the manuscript. T.S., J.S.S., and H-C.C. performed experiments. R.A.M. provided tissues from young and old mice and provided comments on the manuscript. H.A. supervised the project.

## References

Bartzokis, G., Lu, P.H., Tishler, T.A., Peters, D.G., Kosenko, A., Barrall, K.A., Finn, J.P., Villablanca, P., Laub, G., Altshuler, L.L., et al. (2010). Prevalent iron metabolism gene variants associated with increased brain ferritin iron in healthy older men. J Alzheimers Dis 20, 333–341.

Bartzokis, G., Mintz, J., Sultzer, D., Marx, P., Herzberg, J.S., Phelan, C.K., and Marder, S.R. (1994a). In vivo MR evaluation of age-related increases in brain iron. AJNR Am J Neuroradiol 15, 1129–1138.

Bartzokis, G., Sultzer, D., Mintz, J., Holt, L.E., Marx, P., Phelan, C.K., and Marder, S.R. (1994b). In vivo evaluation of brain iron in Alzheimer’s disease and normal subjects using MRI. Biological psychiatry 35, 480–487.

Bayeva, M., Khechaduri, A., Puig, S., Chang, H.C., Patial, S., Blackshear, P.J., and Ardehali, H. (2012). mTOR Regulates Cellular Iron Homeostasis through Tristetraprolin. Cell metabolism 16, 645–657.

Benkovic, S.A., and Connor, J.R. (1993). Ferritin, transferrin, and iron in selected regions of the adult and aged rat brain. J Comp Neurol 338, 97–113.

Berg, D., Gerlach, M., Youdim, M.B., Double, K.L., Zecca, L., Riederer, P., and Becker, G. (2001). Brain iron pathways and their relevance to Parkinson’s disease. J Neurochem 79, 225–236.

Chang, H.C., Shapiro, J.S., Jiang, X., Senyei, G., Sato, T., Geier, J., Sawicki, K.T., and Ardehali, H. (2021). Augmenter of liver regeneration regulates cellular iron homeostasis by modulating mitochondrial transport of ATP-binding cassette B8. Elife 10.

Chang, H.C., Wu, R., Shang, M., Sato, T., Chen, C., Shapiro, J.S., Liu, T., Thakur, A., Sawicki, K.T., Prasad, S.V., et al. (2016). Reduction in mitochondrial iron alleviates cardiac damage during injury. EMBO Mol Med 8, 247–267.

Daugherty, A., and Raz, N. (2013). Age-related differences in iron content of subcortical nuclei observed in vivo: a meta-analysis. Neuroimage 70, 113–121.

Hahn, P., Song, Y., Ying, G.S., He, X., Beard, J., and Dunaief, J.L. (2009). Age-dependent and gender-specific changes in mouse tissue iron by strain. Exp Gerontol 44, 594–600.

Kawabata, H., Yang, R., Hirama, T., Vuong, P.T., Kawano, S., Gombart, A.F., and Koeffler, H.P. (1999). Molecular cloning of transferrin receptor 2. A new member of the transferrin receptor-like family. J Biol Chem 274, 20826–20832.

Koleini, N., Shapiro, J.S., Geier, J., and Ardehali, H. (2021). Ironing out mechanisms of iron homeostasis and disorders of iron deficiency. J Clin Invest 131.

Lakhal-Littleton, S., Wolna, M., Chung, Y.J., Christian, H.C., Heather, L.C., Brescia, M., Ball, V., Diaz, R., Santos, A., Biggs, D., et al. (2016). An essential cell-autonomous role for hepcidin in cardiac iron homeostasis. Elife 5.

Lei, P., Ayton, S., Finkelstein, D.I., Spoerri, L., Ciccotosto, G.D., Wright, D.K., Wong, B.X., Adlard, P.A., Cherny, R.A., Lam, L.Q., et al. (2012). Tau deficiency induces parkinsonism with dementia by impairing APP-mediated iron export. Nat Med 18, 291–295.

Liang, T., Qian, Z.M., Mu, M.D., Yung, W.H., and Ke, Y. (2020). Brain Hepcidin Suppresses Major Pathologies in Experimental Parkinsonism. iScience 23, 101284.

Lu, L.N., Qian, Z.M., Wu, K.C., Yung, W.H., and Ke, Y. (2017). Expression of Iron Transporters and Pathological Hallmarks of Parkinson’s and Alzheimer’s Diseases in the Brain of Young, Adult, and Aged Rats. Mol Neurobiol 54, 5213–5224.

Noster, J., Persicke, M., Chao, T.C., Krone, L., Heppner, B., Hensel, M., and Hansmeier, N. (2019). Impact of ROS-Induced Damage of TCA Cycle Enzymes on Metabolism and Virulence of Salmonella enterica serovar Typhimurium. Front Microbiol 10, 762.

Qian, Z.M., and Wang, Q. (1998). Expression of iron transport proteins and excessive iron accumulation in the brain in neurodegenerative disorders. Brain Res Brain Res Rev 27, 257–267.

Qiao, B., Sugianto, P., Fung, E., Del-Castillo-Rueda, A., Moran-Jimenez, M.J., Ganz, T., and Nemeth, E. (2012). Hepcidin-induced endocytosis of ferroportin is dependent on ferroportin ubiquitination. Cell Metab 15, 918–924.

Roskams, A.J., and Connor, J.R. (1994). Iron, transferrin, and ferritin in the rat brain during development and aging. J Neurochem 63, 709–716.

Sato, T., Chang, H.C., Bayeva, M., Shapiro, J.S., Ramos-Alonso, L., Kouzu, H., Jiang, X., Liu, T., Yar, S., Sawicki, K.T., et al. (2018). mRNA-binding protein tristetraprolin is essential for cardiac response to iron deficiency by regulating mitochondrial function. Proc Natl Acad Sci U S A 115, E6291–E6300.

Sawicki, K.T., Chang, H.C., Shapiro, J.S., Bayeva, M., De Jesus, A., Finck, B.N., Wertheim, J.A., Blackshear, P.J., and Ardehali, H. (2018). Hepatic tristetraprolin promotes insulin resistance through RNA destabilization of FGF21. JCI insight 3.

Schwartz, A.J., Goyert, J.W., Solanki, S., Kerk, S.A., Chen, B., Castillo, C., Hsu, P.P., Do, B.T., Singhal, R., Dame, M.K., et al. (2021). Hepcidin sequesters iron to sustain nucleotide metabolism and mitochondrial function in colorectal cancer epithelial cells. Nat Metab.

Silvestri, L., Nai, A., Pagani, A., and Camaschella, C. (2014). The extrahepatic role of TFR2 in iron homeostasis. Front Pharmacol 5, 93.

Smith, M.A., Harris, P.L., Sayre, L.M., and Perry, G. (1997). Iron accumulation in Alzheimer disease is a source of redox-generated free radicals. Proc Natl Acad Sci U S A 94, 9866–9868.

Sun, Y., Pham, A.N., and Waite, T.D. (2018). Mechanism Underlying the Effectiveness of Deferiprone in Alleviating Parkinson’s Disease Symptoms. ACS Chem Neurosci 9, 1118–1127.

Wang, J., Chen, G., Muckenthaler, M., Galy, B., Hentze, M.W., and Pantopoulos, K. (2004). Iron-mediated degradation of IRP2, an unexpected pathway involving a 2-oxoglutarate-dependent oxygenase activity. Mol Cell Biol 24, 954–965.

Xiong, X.Y., Liu, L., Wang, F.X., Yang, Y.R., Hao, J.W., Wang, P.F., Zhong, Q., Zhou, K., Xiong, A., Zhu, W.Y., et al. (2016). Toll-Like Receptor 4/MyD88-Mediated Signaling of Hepcidin Expression Causing Brain Iron Accumulation, Oxidative Injury, and Cognitive Impairment After Intracerebral Hemorrhage. Circulation 134, 1025–1038.

Xu, J., Jia, Z., Knutson, M.D., and Leeuwenburgh, C. (2012). Impaired iron status in aging research. Int J Mol Sci 13, 2368–2386.

Zecca, L., Gallorini, M., Schunemann, V., Trautwein, A.X., Gerlach, M., Riederer, P., Vezzoni, P., and Tampellini, D. (2001). Iron, neuromelanin and ferritin content in the substantia nigra of normal subjects at different ages: consequences for iron storage and neurodegenerative processes. J Neurochem 76, 1766–1773.

Zhang, X., Gou, Y.J., Zhang, Y., Li, J., Han, K., Xu, Y., Li, H., You, L.H., Yu, P., Chang, Y.Z., et al. (2020). Hepcidin overexpression in astrocytes alters brain iron metabolism and protects against amyloid-beta induced brain damage in mice. Cell Death Discov 6, 113.

